# Activated carbon ameliorates type 2 diabetes via metabolic remodeling of the gut microbiota

**DOI:** 10.1101/2024.10.15.618493

**Authors:** Cai-Xia Zhao, Yu-Tong Wu, Xiao-Yu Guo, Yin Wang, Jian-Qiang Su

## Abstract

Type 2 diabetes (T2D) is a major public health concern worldwide and there has been increasing attention on the role of natural dietary drugs in diabetes therapy. However, the effects of these drugs on gut microbial composition, functional potentials and metabolisms remain unclear. Here, we conducted integrated 16S rRNA sequencing and metabolomic analyses in T2D GK rats and healthy Wistar rats that exposed to four natural dietary drugs (highly porous activated carbon, wheatgrass, dandelion and corn stigma). Oral administration of activated carbon and dandelion decreased the body weight gain in both high-fat diet (HFD) GK rats and Wistar rats. Significantly lower level of blood glucose was observed in GK rats with activated carbon intervention. A group of beneficial bacteria and metabolites were promoted, and the endotoxin-producing bacteria were inhibited by dietary drugs, especially for the activated carbon diet. Oral administration of activated carbon resulted in metabolic changes and anti-inflammatory effects that decreased both high-fat diet-induced obesity and diabetes. The beneficial effects of increased positive responders are related to improved carbohydrate and amino acid metabolism, regulated inflammatory mediators, with simultaneous reduction of detrimental compounds such as lipopolysaccharide (LPS) synthesis and modification of the gut microbiome. These findings highlight the effectiveness of natural dietary drugs, with a particular emphasis on activated carbon, and establish a foundation for tailoring the use of these drugs in T2D therapy.

**Importance:** Our findings highlight the significant hypoglycemic effect of activated carbon, demonstrating its potential to remodel the gut microbiota, improve carbohydrate and amino acid metabolism, regulate inflammatory mediators, and reduce detrimental compounds such as lipopolysaccharide (LPS). These results suggest that dietary intervention with activated carbon could be a noninvasive and accessible method for improving diabetes management, providing novel insights into the role of natural dietary drugs in metabolic health and diabetes therapy.

## Introduction

Diabetes mellitus are one of the major public health concerns worldwide, and their prevalence has continued to increase over the past half-century. The incidence of diabetes has reached a staggering 463 million individuals in 2019, as claimed by the International Diabetes Federation (IDF). The IDF estimates that there will be 578 million adults with diabetes by 2030 and 700 million by 2045, with anticipated healthcare costs of around $850 billion annually (1). This epidemic is mainly due to the increase in the Type 2 diabetes (T2D) incidences, a global epidemic endocrine disease characterized by insulin resistance and β-cell dysfunction. The etiology of T2D is multifactorial, consisting of genetic susceptibility and environmental factors, including lifestyle, medical conditions, dietary components, and gut microbial alterations (2–4).

There are mounting evidences indicate that gut microbiome signatures distinguish T2D patients from healthy individuals (5–7), suggesting the essential roles of gut microbes in managing T2D and maintaining host health, which can be partially attributed to the production of microbial-derived short-chain fatty acids (SCFAs) (8). Previous large-scale T2D-related metagenomics research in China has shown increased abundances of opportunistically pathogenic genera *Clostridium* and decreased abundances of butyrate-producing genera *Roseburia*, *Faecalibacterium*, and *Eubacterium* associated with T2D patients (9). Zhao et al. (10) found a selected group of SCFA-producing strains were promoted while producers of detrimental metabolites were diminished following the uptake of dietary fibers in T2D patients. In addition, Hosomi et al. (11) found that oral administration of *Blautia wexlerae* to mice decreased both high-fat diet-induced obesity and diabetes. Dietary supplementation of corn bran arabinoxylan to adults modulates both gut microbiome composition and the output of health benefic SCFAs (12). As a result, there is a burgeoning interest towards modifying the microbiome through lifestyle and dietary changes and/or probiotic administration to help alleviating and preventing various disease risks associated with T2D.

Oral synthetic antidiabetic agents have been broadly adopted as therapies for the treatment of T2D, while some traditional natural herbal remedies have been reported to be a new kind of compounds that are more effective, less costly and less toxic as hypoglycemic agents. Traditional natural herbal medicines for diabetes have demonstrated potential to alleviate diabetic symptoms, enabling recovery and improving health. Wheatgrass (*Triticum aestivum* L.), dandelion (*Taraxacum Officinale*) and corn stigma (*Stigma Maydis*) are herbs with remarkable applications in this respect. Wheatgrass, young germinated shoots of *Triticum aestivum* L., has been performed to exhibit anti-oxidant and anti-diabetic therapeutic effects (13,14). Studies have pointed out that dandelion supplements can be advantageous in preventing diabetic complications as an antioxidant therapy (15,16). Corn stigma, a waste product of corn cultivation, has been proven to have benefits in the management of infections, obesity and diabetes (17). Activated carbon with highly porous, an FDA-approved product for using in humans and generally regarded as safe, has been reported to improve the therapeutic efficacy of common drugs using murine models of corneal and genital herpes infections (18). However, there is dearth of information on data regarding the availability and mechanism of these natural drugs in vivo anti-diabetic activities. The comprehensive profiling of microbiota, functional pathways and metabolic dysbiosis in T2D after these natural drugs intervention is lacking, and the dynamic change of body weight and fasting blood glucose is required for further identification.

To decipher the effect and mechanism of the intervention of traditional natural herbal medicines, we performed integrated 16S rRNA sequencing and metabolomic analyses in two independent cohorts: one with spontaneously T2D GK rats and another with healthy Wistar rats. We identified microbial species and fecal metabolites that might be related to T2D risk. Our results showed that activated carbon could regulate the structure and diversity of intestinal microflora, change the relationships between the gut microbiota and metabolites, thus improving gut dysbiosis and clinical performance. This further emphasizes the understanding that activated carbon might possess the potentiality as an effective natural functional ingredient in the management of diabetes.

## Results

### Body weight and blood glucose levels

Rats fed with a high-fat diet (HFD) (GK rats) resulted in significant weight gain compared to rats with a standard normal diet (Wistar rats). However, oral administration of activated carbon and dandelion during HFD feeding concurrent with decreased body weight gain, compared with non-supplemented these two drugs within GK rats (*P* < 0.05). Consistent results were also observed within Wistar rats under these two drugs intervention (Fig. 1A). These results suggest that activated carbon and dandelion has potential to contribute to the prevention of obesity.

**Fig. 1.**
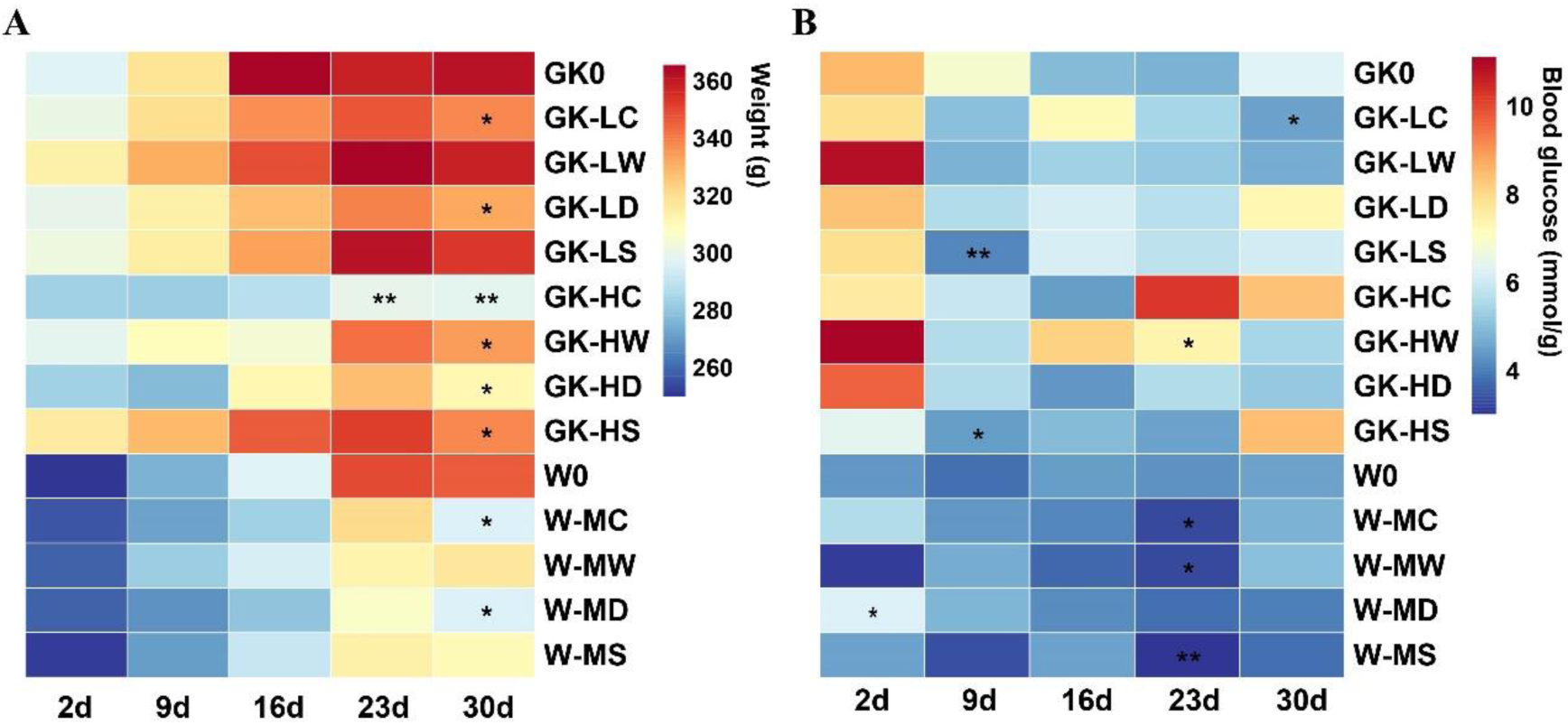
Dynamic changes of body weight and fasting blood glucose (FBG) in GK and Wistar rats during 30 days. (A) Body weight; (B) FBG. *P < 0.05, **P < 0.01 using one-way ANOVA test or Kruskal-Wallis tests for intra-group comparisons. Data in the figures were presented as mean of each group (n = 4).

Compared with Wistar rats, GK rats showed higher blood levels of glucose under fasting conditions. Rats that received low dose activated carbon intervention had reversed this trend (*P* < 0.05). Significant difference in the blood glucose levels of W groups was not observed, except that W-LD achieved a significant reduction by day 23 (Fig. 1B).

### Gut microbiota

To examine the variation in microbial communities among different treatments, 16S rRNA gene sequencing was conducted on fecal samples collected at the last sampling time point (day 30). The Wistar groups showed a tendency towards higher species diversity, while the species richness of the Wistar groups was comparable to the high dose treatment in GK groups and lower than that in the low dose groups. Although there was an effect of increased alpha diversity in GK-LC and decreased alpha diversity in GK-HW when compared with GK0, the bacterial profile remained relatively similar to the GK0 (Fig. 2A and Fig. 2B). Principal coordinate analysis (PCoA) based on weighted UniFrac distance metrics of gut bacterial communities indicated that samples were separated by dietary drug treatment. The bacterial community profiles in the activated carbon treatments indicated a significant divergence, with increased and decreased compositional dissimilarity in the W-MC group and activated carbon treated GK groups, respectively (Fig. 2C, Fig. S2A, Fig. S3A). This suggests a more and less heterogeneous community structure among these three groups than in their corresponding controls. A similar pattern was observed in all high dose treatments and GK-LS group (Fig. S2B to S2D, Fig. 2F).

**Fig. 2.**
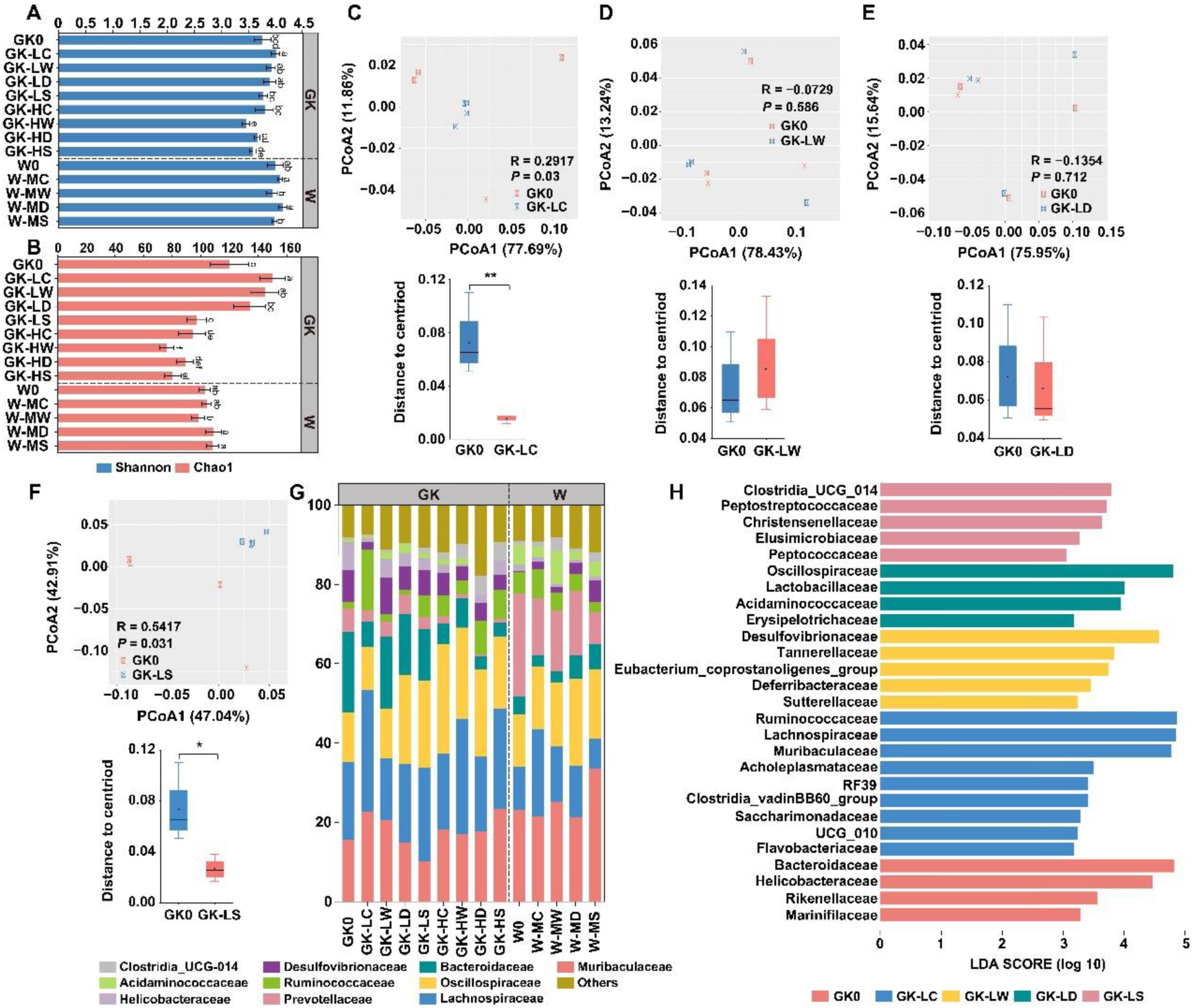
The structural shift of gut microbiota in GK and Wistar rats. (A) The microbial community diversity (Shannon index) and (B) richness (Chao 1 index). (C), (D), (E), (F) Principal coordinate analysis (PCoA) based on weighted-UniFrac distance and inner-group dissimilarity in GK groups under low dose dietary drug intervention. (G) The relative abundance of microbial taxa at family level, family with a relative abundance < 1% in each sample are merged into RA < 1%. (H) Differences in bacterial taxonomy at the family level were ranked according to the LefSe analysis.

Despite this, there were concomitant pronounced relative abundance variations within each group across different drug treatments. Each diet resulted in a unique set of differential taxa between the microbial communities, including variations in phyla (Fig. 3A). It’s worth to note that rats administrated with activated carbon from either the GK or W groups consistently showed higher abundance of Firmicutes and lower abundance of Bacteroidetes than those in the other drug administration groups (Fig. 3B to 3E). Notably, the ratio of Firmicutes to Bacteroidetes (F/B) significantly increased in all groups treated with activated carbon (*P* < 0.05) (Fig. 3F). The majority of the taxa at the family level are derived from Muribaculaceae (10.21-33.57%), Lachnospiraceae (7.63-30.71%) and Oscillospiraceae (10.67-27.68%) (Fig. 2G).

**Fig. 3.**
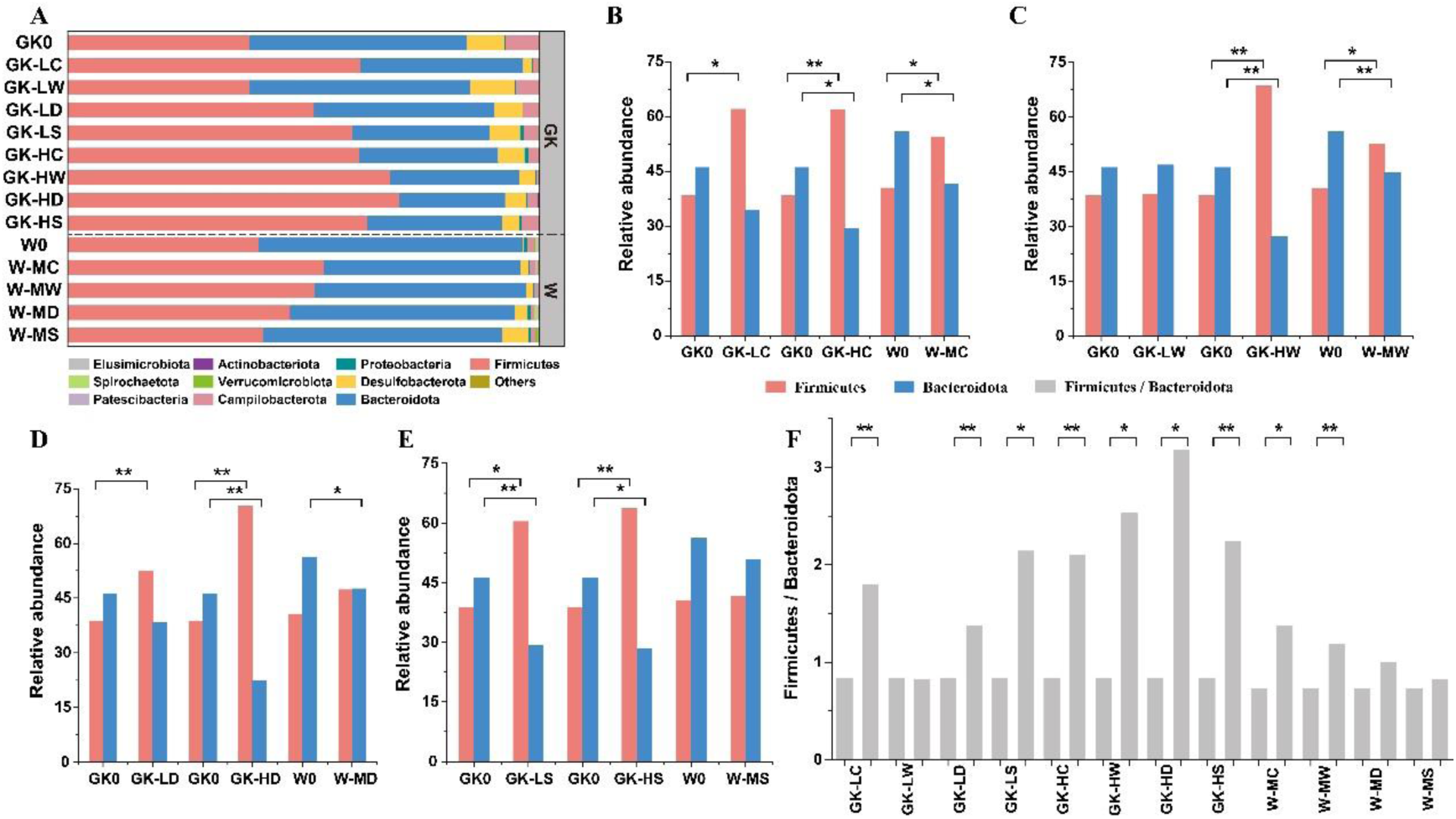
The relative abundance of microbial taxa at phylum levels among engrafted different drugs. (A) The relative abundance (RA) of the top 10 phyla. "Others" is used to denote the remaining phyla. (B) RA of Firmicutes and Bacteroidetes for low (GK1), medium (W1) and high (GK5) dose of activated carbon intervention. (C) RA of Firmicutes and Bacteroidetes for low (GK2), medium (W2) and high (GK6) dose of wheatgrass intervention. (D) RA of Firmicutes and Bacteroidetes for low (GK3), medium (W3) and high (GK7) dose of dandelion intervention. (E) RA of Firmicutes and Bacteroidetes for low (GK4), medium (W4) and high (GK8) dose of Corn Stigma intervention. (F) The individual effect of dietary drugs on Firmicutes to Bacteroidetes ratio compared to the respective control group. \**P* < 0.05, \*\**P* < 0.01.

Moreover, the changes in communities were similar in response to the same drugs, indicating that related taxa respond similarly to a particular diet and that different drugs have distinct effects on the relative abundance of the taxa. We used linear discriminant analysis effect size (LefSe) to provide a ranked list of intestinal bacteria that differed between control group and post-intervention microbiomes for each dietary drug. Low dose activated carbon diet significantly elevated the relative abundance of some potential beneficial bacteria such as Ruminococcaceae, Lachnospiraceae, Muribaculaceae and Saccharimonadaceae (*P* < 0.05) (Fig 2H). Meanwhile, Bacteroideaceae and Desulfovibrionaceae exhibited decreased abundance to varying degrees (Fig 2G). After increasing the dosage, we found significantly elevated levels of Oscillospiraceae, Rikenellaceae and Izemoplasmatales, while Prevotellaceae declined in abundance (*P* < 0.05) (Fig. 2G, Fig. S2E). Medium dose activated carbon-fed healthy subjects displayed increased abundance of Ruminococcaceae, Lachnospiraceae, Rikenellaceae and Izemoplasmatales, and decreased abundance of Bacteroideaceae (Fig. 2G, Fig. S3E).

In the low dose wheatgrass diet, the abundance of Eubacterium_coprostanolinenes_group, Desulfovibrionaceae and Deferribacteraceae increased, accompanied by decreased abundance of Helicobacteraceae and Lachnospiraceae (Fig. 2G, Fig. 2H). A higher abundance of Lachnospiraceae, as well as lower levels of Bacteroideaceae, Helicobacteraceae, Desulfovibrionaceae and Prevotellaceae were observed in high dose wheatgrass treated group (Fig. 2G, Fig. S2E). In medium dose W-LW group, increased abundance of Clostridia_UCG_014 and Eubacterium_coprostanolinenes_group, as well as the decreased abundance of Helicobacteraceae was observed (Fig. 2G, Fig. S3E). As to the dandelion intervention groups, abundance of Oscillospiraceae and Lactobacillaceae was increased in low dose treated group (Fig. 2H). Significantly elevated levels of Ruminococcaceae and Clostridia_UCG_014, Lactobacillaceae, Eubacterium_coprostanolinenes_group, Bifidobacteriaceae as well as Defluviitaleaceae was observed in high dose treated group and abundance of Christensenellaceae was increased in medium dose treated group (*P* < 0.05) (Fig. S2E, Fig. S3E). The abundance of Helicobacteraceae and Prevotellaceae all decreased to varying degrees under the different dose treatment of the dandelion (Fig. 2G). Families related to Clostridia_UCG_014 and Christensenellaceae became enriched in low dose corn stigma fed rats, while families related to Muribaculaceae and Enterobacteriaceae were enriched in high dose fed rats (Fig. 2H, Fig. S2E). In contrast, the proportion of Muribaculaceae, Helicobacteraceae and Prevotellaceae were decreased to varying drgrees under these interventions (Fig. 2G). Furthermore, the abundance of Akkermansiaceae, Lactobacillaceae, Bacteroidaceae and Desulfovibrionaceae was increased in healthy rats given the medium dose treatment. And the level of potential beneficial bacteria Ruminococcaceae and Lachnospiraceae decreased (Fig. 2G, Fig. S3E).

Some families that showed diet-specific differences between communities were only detected in one of the communities, such as Saccharimonadaceae and Izemoplasmatales, which were exclusively detected in the activated carbon diet. These results provide clear evidence of the dominant role of diet in shaping the microbial composition of the gut, as well as the specific compositional effects of activated carbon in relation to other drug treatments.

Additionally, we utilized PICRUSt2 to predict the functional features of the gut microbiome using 16S rRNA gene data. A PCoA plot based on the Bray Curtis distance of the KEGG orthologs (KOs) showed a significant difference in the activated carbon treatment groups, indicating distinct functional potentials of the gut microbiomes. Intervention-specific effects was observed among the low dose treatment GK groups and the Wistar groups. For instance, fructose and mannose metabolism, glycolysis/gluconeogenesis, starch and sucrose metabolism and the pentosephosphatate pathway were observed only in the activated carbon treated groups. At high dose treatment levels, the carbohydrate metabolism pathway was enriched in all treated groups except for the activated carbon group. It is worth noting that lipopolysaccharide (LPS) synthesis was reduced in all treatment groups (Fig. 4, Fig. S2F, Fig. S3F).

**Fig. 4.**
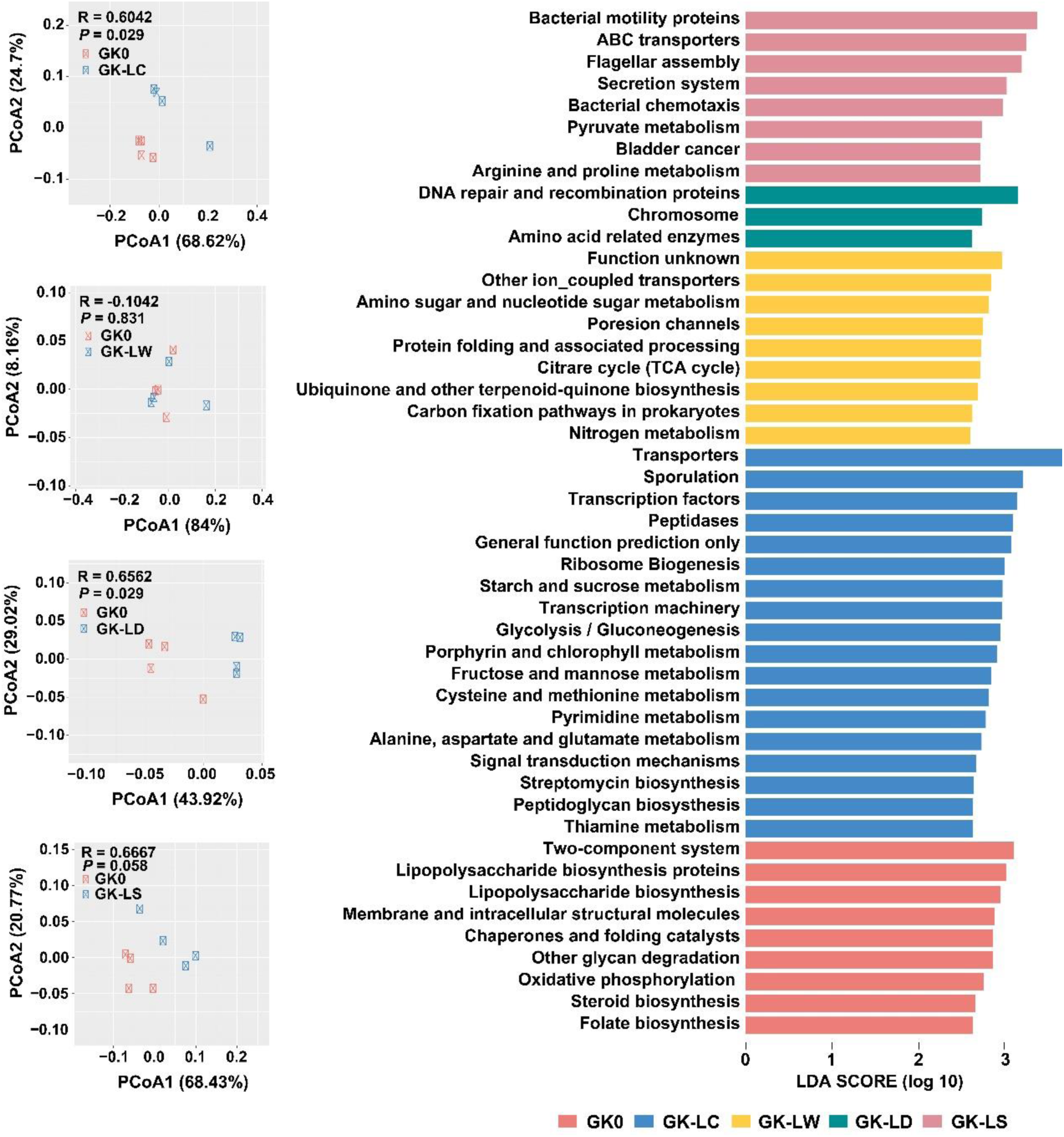
Functional analysis reveal functions in GK groups under low dose dietary drug intervention. The PCoA plot based on the Bray-Curtis distance of KOs by ANOVA are shown on the left, while the differences in predicted metabolic functions are shown on the right (*P* < 0.05 and LDA > 2.5).

### Metabolite profiles

To further investigate the effects of dietary drug interventions on changes in the gut microbiome on intestine’s metabolic profile, untargeted metabolomics of fecal samples was performed. In total, 4751 and 4036 mass spectra peaks were detected in positive and negative ion modes, respectively, with a total of 919 and 347 metabolites were annotated. PLS-DA showed substantial overlap of GK groups, except for GK-LC and GK-HC groups. Similarity, we found that a large fraction of these metabolites also showed a significant diet effect in the Wistar groups, and the metabolite profiles of the W-MC group were distinct from the others, indicating that the activated carbon treated groups have a dissimilar metabolic profiles compared to the control (Fig. 5A).

**Fig. 5.**
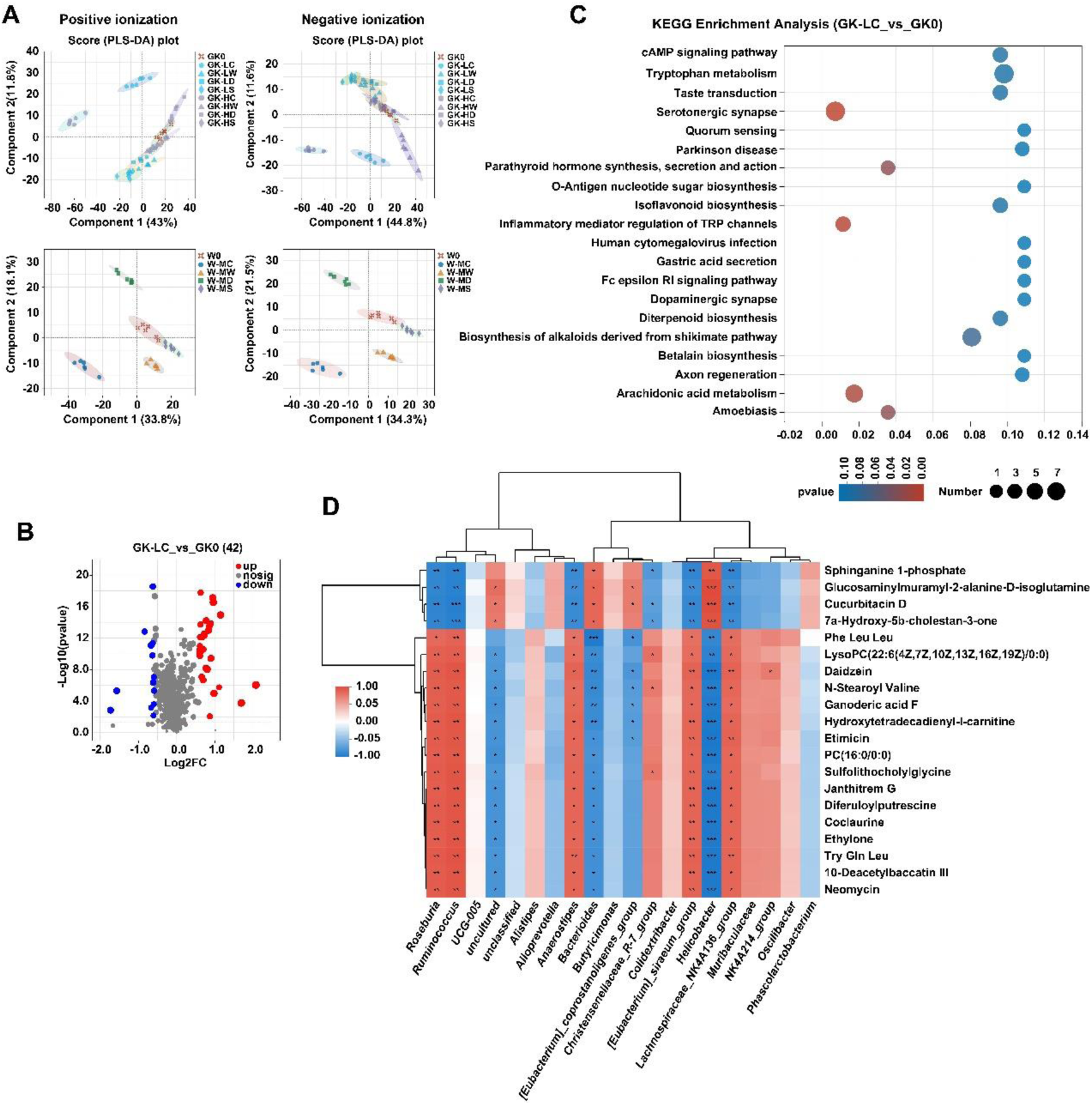
Fecal metabolic patterns. (A) Partial least squares discriminant analysis (PLS-DA) of positive and negative ionization dataset for GK and Wistar rats. (B) Volcano maps of differential metabolites in the low dose activated carbon treated GK1 group. Significantly upregulated and downregulated genes are shown in red and blue, respectively. Genes with no significant changes (nosig) in transcription are shown in grey. (C) KEGG pathway enrichment analysis between the GK0 and GK1 group. (D) Heatmap analysis of the Spearman’s correlation between 20 differential metabolites and 20 key bacteria genera. The red squares indicate positive correlations, whereas the blue squares indicate negative correlations. The metabolite clustering tree is shown on the left. The distance between branches shows the closeness in the expression pattern of metabolites.

Compared to GK0 and W0, there were 42, 10, 6, 16, 64, 8, 15, 6 different metabolites in GK-LC to GK-HS and 31, 6, 5, 5 different metabolites in W-MC to W-MS, respectively (Supplementary Data 1 to 12). The proportion of significantly changed metabolites in activated carbon treated rats were the highest compared to the other treatments (Fig. 5B, Fig. S4A, Fig. S5A, Fig. S6A, Fig. S7A). Specifically, when comparing unique metabolites in activated carbon treated groups and other groups in GK and Wistar rats, a total of 8 (with 4 increased and 4 decreased) and 27 (with 6 increased and 21 decreased) significantly different metabolites were identified, respectively (*P* < 0.05).

According to HMDB classification, these discriminant metabolites belonged to 5 superclass taxonomy, mainly related to phenylpropanoids and polyketides, lipid and lipid-like molecules, organic acids and derivatives (Supplementary Data 13 and 14). Inspection of the differentially abundant metabolic pathways within each diet revealed diet-specific changes in metabolites. According to KEGG pathway enrichment analysis, there was a significant over-representation of metabolites in the low dose activated carbon treatment GK group related to serotonergic synapse, inflammatory mediator regulation of TRP channels, and arachidonic acid metabolism (Fig. 5C) (*P* < 0.05). Tryptophan metabolism and steroid hormone biosynthesis were also significantly enriched after increasing the treatment dose. More discriminant metabolic pathways were enriched in the activated carbon treatment Wistar group: steroid biosynthesis, isoflavonoid biosynthesis, flavonoid biosynthesis, aldosterone synthesis, and secretion (Fig. S4B) (*P* < 0.05). However, these metabolic pathways were rarely enriched in the other treatment groups (Fig. S5, Fig. S6, Fig. S7). These results suggest that dietary drug treatment may be accompanied by changes in metabolites. Activated carbon groups may regulate the host metabolism by significantly increasing the expression level of functional metabolites.

### Correlation analysis between gut microbiota and fecal metabolites

To investigate the microbiota–metabolite correlations between treated groups, Procrustes analysis was performed. The results highlighted a significant association between the microbiota and metabolites (Fig. S8). Pearson correlation analysis between gut microbiota and differential metabolites based on the top 20 significant differential genera and metabolites showed that the genera affiliated to Firmicutes were the most connected within each group, followed by those from Bacteroidota and Campilobacterota.

Regarding the GK-LC group (low dose treatment of activated carbon), we observed that the genera *Roseburia*, *Ruminococcus*, *Anaerostipes*, *Christensenellaceae_R-7_group*, *[Eubacterium]_siraeum_group*, *Lachnospiraceae_NIK4A136_group* and *NIK4A214_group* were negatively correlated with downregulated metabolites of the Sphingolipids biosynthesis (Sphinganine 1-phosphate), Carboxylic acids and derivatives biosynthesis (7a-Hydroxy-5b-cholestan-3-one), and Steroids and steroid derivatives biosynthesis (Glucosaminylmuramyl-2-alanine-D-isoglutamine and Cucurbitacin D). These genera were also positively associated with the biosynthesis of some upregulated metabolites, such as Daidzein, LysoPC (22:6(4Z,7Z,10Z,13Z,16Z,19Z)/0:0), PC (16:0/0:0)), Ganoderic acid F, 10-Deacetylbaccatin III, N-Stertoyl Valine, Tyr Gln Leu and Diferuloylputrescine, further indicating a synergistic relationship between these taxa and metabolites (Fig. 5D). Interestingly, *Roseburia*, *Ruminococcus*, *Anaerostipes*, and *[Eubacterium]_siraeum_group*, along with *Lachnospiraceae_NIK4A136_group*, all of which belong to Lachnospiraceae and Ruminococcaceae, showed significantly increased abundance (Fig. 2G). However, the abundance of *Bacteroides* and *Helicobacter*, which exhibit opposite correlations with the metabolites mentioned above, were significantly reduced (*P* < 0.05) (Supplementary Data 15). After being treated with high dose activated carbon (GK-HC), several genera including *Ruminococcus*, *UCG-005*, *Alistipes*, *Clostridia_UCG-014*, *Lactobacillus*, *Muribaculaceae*, and *NIK4A214_group* exhibited a strong positive correlation with upregulated Isoflavonoids biosynthesis (Daidzein), Carboxylic acids and derivatives biosynthesis (N-Stearoyl Valine), and Cinnamic acids and derivatives biosynthesis (Diferuloylputrescine) (Fig. S4C). The abundance of these genera, except for *Muribaculaceae*, were significantly elevated (*P* < 0.05). Noticeably, *Rikenellaceae_RC9_gut_group*, *Bacteroides*, *Butyricimonas* and *Helicobacter*, were detected with reduced abundance, and showed negative correlation with the metabolites as compared to the aforementioned genera (Supplementary Data 15). Furthermore, the medium dose activated carbon supplement (W-MC) showed induced abundance in *Ruminococcus*, *UCG-005*and *NIK4A214_group*, and were positively correlated with upregulated Glycitein, Genistein, Apigenin, VASICINONE, DG (PGF1alpha/2:0/0:0), and N1, N10-Dicoumaroylspermidine. Additionally, some steroids-related synthesis (e.g., Drostanolone, 4alpha-Formyl-5alpha-cholesta-8-en-3beta-ol and 16a-Hydroxydehydroisoandrosterone), Atenolol and 6-O-Acetylaustroinulin were significantly inhibited (*P* < 0.05) (Fig. S4C). Meanwhile, the abundance of the genera exhibiting opposite correlations with the relevant metabolites, such as *Christensenellaceae_R-7_group*, *Helicobacter*, *Lachnospiraceae_NK4A136_group* and *Prevotellaceae_NK3B31_group*, were significantly reduced (*P* < 0.05) (Supplementary Data 16).

There was a significant positive correlation between the 3,7,11,15,23-Pentaoxolanost-8-en-26-oic acid, Arenaine, Sterebin A, VINCAMINE biosynthesis and the genera *Anaerostips*, *Butyricimonas*, *Helicobacter* under the treatment of low dose wheatgrass, but the expression of these metabolites decreased significantly (*P* < 0.05) (Fig. S5A). In the high dose treatment, more genera, e.g., *Alloprevotella*, *Bacteroids*, and *Lachnospiraceae_NK4A136_group*, were negatively correlated with downregulated Piperidines biosynthesis ((+/-)-Pelletierine), while the abundance of these genera significantly decreased except *Alloprevotella* and *Anaerostipes* (*P* < 0.05) (Fig. S5B, Supplementary Data 15). In the W-MW group, the abundances of *Ruminococcaceae*, *[Eubacterium]_coprostanoligenes_group* and *Clostridia_UCG-014* significantly increased (*P* < 0.05), which were negatively correlated with Uridine-5’-Monophosphate and Eptastigmine biosynthesis, but positively correlated with Asn Pro Leu, Phe Leu Leu and MG (PGE2/0:0/0:0) (Fig. S5C, Supplementary Data 16).

Under low dose treatment of dandelion, rising *Romboutsia*, *UCG-005*, *Lactobacillus* and *Phascolarctobacterium* were positively assoiciated with upregulated isoflavonids (Luteolin). Notably, we observed an increased correlation between the genera and metabolites under the high dose treatment. *Ruminococcaceae*, *Ruminococcus*, *UCG-005*, *Christensenellaceae_R-7_group*, *Clostridia_UCG-014*, *Clostridium_sensu_stricto_1*, *Lactobacillus and NIK4A214_group* with significantly increased abundance were positively correlated with upregulated metabolites, such as 6-isobutyl-4-hydroxy-2-pyrone, Diferuloylputrescine and Luteolin (*P* < 0.05). In the W-MD group, *[Eubacterium]_coprostanoligenes_group*, *Helicobacter*and *Phascolarctobacterium* were positively correlated with 1-Palmitoylphosphatidylcholine, N1, N10-Dicoumaroylspermidine and Gonyautoxin I synthesis, while they are also inhibited the purine nucleotides pathway. But the abundance of *Helicobacter* and *[Eubacterium]_coprostanoligenes_group* was significantly decreased (Fig. S6C, Supplementary Data 15 and 16).

In T2D rats with under corn stigma treatment, a significant positive correlation was observed between the rising *Anaerostipes*, declining *Helicobacter* and *Lachnospiraceae_NIK4A136_group*, as well as the downregulated metabolites: 3,7,11,15,23-Pentaoxolanost-8-en-26-oic acid, (3S,5R,6R,7E)-3,5,6-Trihydroxy-7-megastigmen-9-one and 7-Amino-4-(trifluoromethyl) coumarin biosynthesis. The abundance of genera that were negatively correlated with the same metabolites significantly increased (*P* < 0.05). Upon increasing the medication dosage, the abundance of genera *Bacteroides* and *Butyricimonas* decreased and were correlated with the downregulated 3,7,11,15,23-Pentaoxolanost-8-en-26-oic acid. In the W4 group, *Clostridia_UCG-014*, *Lactobacillus* and *Muribaculaceae* were detected with increased abundance. *NIK4A214_group* were positively correlated with upregulated Asn Pro Leu, Cholylglutamic acid, Phe Leu Thr and 3-Amino-4-methoxybenzanilide (Fig. S7C, Supplementary Data 15 and 16).

The interrelationships between metabolites and genera indicate the potential involvement of gut microbiota in regulating host metabolism. However, the correlations observed between metabolites and changes in the gut microbiome may differ based on the treatment and dose. These findings proved that the consumption of dietary drugs elicits individual responses on fecal metabolites that have the ability to reshape the structure of the gut microbiota.

## Discussion

In this study, our multi-omics analyses and animal experiments firstly deciphered the impact of a 30-day, different-dose and -type natural dietary drug administration on the composition and function of the fecal bacterial community in T2D GK rats and healthy Wistar rats. We explored the relationship and underlying mechanism between the gut microbiota and metabolism.

In our study, the Wistar groups had lower species richness compared to the low dose treatment GK groups, despite significant clinical improvements. However, species richness tended to be similar to the Wistar groups in the high dose treatment groups. This may be related to the ratio of Bacteroidetes/Firmicutes, as Firmicutes is a highly diverse division, which is in agreement with the findings of Larsen et al. (19). The gut microbiota compositions in drug diet rats were dominantly characterized by Firmicutes and Bacteroidaceae, which is consistent with most previous studies. Besides, we observed a significantly higher Firmicutes/Bacteroidaceae ratio, as reported by Ross et al. (5). Previous studies have also reported that this ratio is significantly correlated with glucose metabolism and insulin sensitivity (19–21), which is consistent with our observations in the activated carbon diet. At the family level, we found that potential beneficial bacteria, including Lachnospiraceae, Oscillospiraceae, Muribaculaceae, Ruminococcaceae and Izemoplasmatales, were more increased after activated carbon treated, while harmful bacterial, including Bacteroideaceae and Desulfovibrionaceae, were decreased. However, the other three drug diet groups seemed to have lower specificity and promoted multiple deleterious bacteria of Bacteroideaceae, Desulfovibrionaceae and Enterobacteriaceae, as well as Prevotellaceae, which were positively correlated with diabetes and impaired glucose tolerance, as reported previously (22). It is noteworthy that all of these four treatments did have some inhibitory effect on the harmful bacteria compared to the control group. Additionally, the results indicate that 30 genera, including SCFA and bile acid-producing genera (e.g., *Alistipes*, *Butyricimonas*, *Lactobacillus*, *Oscillibacter*, *Ruminococcus*, *Romboutsia*, *Muribaculaceae*, *Turicibacter* and *Parabacteroides*), as well as harmful genera (e.g., *Bacteroides* and *Alloprevotella*), are widely related to metabolic disorders such as flavonoids, isoflavonoids, sphingolipids, glycerophospholipids and steroids. Evidences have also shown a strong association between the gut microbiome and metabolites after dietary intervention, particularly short-chain fatty acids (SCFAs) that are produced by certain gut microbiota promoted by dietary fibers (10). This provides clues to find the interaction of each omics and may be a worthwhile target for probiotic manipulation.

Gram-negative bacteria mentioned above, which can produce lipopolysaccharides (LPS), have been considered as potent stimulators of inflammation and can exhibit endotoxaemia (22,23). Our results demonstrate a significant correlation between these bacteria and certain metabolites, such as sphingolipids, which have been strongly associated with adiposity, inflammation and metabolic impairment in previous studies (24–26). In GK rats of the activated carbon diet, we observed a significant decreased in sphingolipids, which was accompanied by decreased body weight and FBG. These findings suggest that the gut microbiome may contribute to the development of metabolic disease or that the microbial community structure may serve as a diagnostic biomarker of metabolic disease onset.

Polyphenol (e.g., flavonoids, isoflavonoids and cinnamic acids), bioactivate components known as "candidates prebiotics", was enriched in the activated carbon diet and largely contributed to the positive association observed between this module and the beneficial bacteria mentioned earlier. This is consistent with previous research that shows dietary polyphenols exert prebiotic-like effects with the growth of beneficial bacteria and the inhibition of pathogenic bacteria (27,28), thus contributing to a healthier gut environment. Molinari et al. reported that the administration of foods rich in polyphenols coincided with a significant inhibition of α-glucosidase and the promotion of the translocation of GLUT4 to the plasma membrane, resulting in a reduced incidence and better metabolic control of T2D in both human and animal epidemiological studies (28,29). In the study of cardiovascular diseases, it was found that individuals with reduced blood pressure, total cholesterol and triglycerides had increased abundance of Lachnospiraceae, Ruminococcaceae and Oscillospirales (30). Our study also found a significant increase in these well-known beneficial bacteria after the intervention of activated carbon, indicating a positive correlation with reducing triglycerides and cholesterol.

Furthermore, the increase of Lachnospiraceae, in particular, has been associated with obesity, with the protective mechanism attributed to higher butyrate production (31). This ability is found in many species within the Lachnospiraceae family, including *Anaerostipes*, *Roseburia* and *Lachnospiraceae_NK4A136_group*. Previous studies have reported a significant increase in beneficial bacteria such as *Roseburia* in subjects with reduced body fat (32), which is similar to our results. These bacteria were also correlated with certain bacteria-originated or bacteria-metabolized metabolites, such as branched-chain amino acids (BCAAs), prenol lipid and glycerophospholipid. BCAAs (e.g., N-Stearoyl Valine and leucine), which may play a crucial role in regulating insulin action and act as a putative biomarker for future T2D incidence (33,34), were enriched in the activated carbon diet compared to other groups. Regarding prenol lipid and glycerophospholipids, which are crucial structural components of cell membranes and key signaling molecules in numerous pathophysiological states, they were also significantly upregulated in the activated carbon diet (35,36). Moreover, Saccharimonadaceae, which belongs to Saccharibacteria, was only present in the activated carbon diet and may be essential for the immune response, oral inflammation and inflammatory bowel disease (37). Izemoplasmatales, related to amino acid biosynthesis, were also enriched in the same diet (38).

Lactobacillus, which belongs to Lactobacillaceae, represents a heterogeneous group with well-documented immune-modulating properties and may potentially contribute to chronic inflammation in diabetic subjects (39). However, we found that high-dose activated carbon group, with more starch added, did not have a significant effect compared to the low-dose group, as excessive starch consumption has been reported to induce metabolic diseases (40). These findings suggest that the supplementation of activated carbon plays a multifactorial role in regulating carbohydrate metabolism, inflammatory mediators and lipid-like molecules in rats.

Altogether, these results demonstrate a link between changes in levels of bacterial taxa differentially represented across the various diabetic drugs and alterations in systemic levels of metabolites and host metabolic phenotypes. The positive responders promoted by dietary drugs are likely the key players in maintaining the mutualistic relationship between the gut microbiota and the host. By promoting this active group, not only a beneficial function enhanced, but a gut environment that may help to regulate the colonization and pathogenicity of detrimental microbes causing obesity and hyperglycemia is also maintained (41,42). This is seen in the fact that gram-negative bacteria were significantly underrepresented in the beneficial genera enriched activated carbon diet groups compared to other groups. These findings are not only helpful for future investigations to develop diagnostic or microbial therapeutic tools for T2D but also relevant for animal studies in the nutrition field.

## Conclusions

In this study, we demonstrated that the activated carbon diet is an effective treatment option to have a hypoglycemic effect by remodeling the gut microbiota and significantly impacts the metabolites of T2D. If efficacious, a dietary intervention with activated carbon could be a well-accepted, noninvasive and highly accessible method for their potential merit in improving diabetes.

## Materials and methods

### Animal design

Animal experiments were conducted in accordance with the guide for the Care and Use of Laboratory Animals and were approved by the Ethics Committee of Institute of Urban Environment (approval number: 2023003). Male spontaneously type 2 diabetes (T2D) GK rats (aged 8 weeks) and non-diabetic male Wistar rats (aged 8 weeks) were obtained from the SPF Cavens Laboratory Animal Co., Ltd. (Changzhou, China) and Shengchang Biotechnology Co., Ltd. (Shanghai, China), respectively. All rats were randomly assigned to either the control or experimental groups and housed under standard environmental conditions (temperature, 23 ± 2 °C; humidity, 55 ± 10%) with a 12-hour light-dark cycle. The GK rats were fed with high-fat D12451 (45 % fat) diet, while the Wistar rats fed with normal diet (GB 14924, 12.11 % fat). All rats were allowed free access to deionized water and diet for one week acclimatization before being randomly divided into nine GK groups: control GK (GK0), GK plus low dose activated carbon (GK-LC), GK plus low dose wheatgrass (GK-LW), GK plus low dose dandelion (GK-LD), GK plus low-dose corn stigma (GK-LS), GK plus high-dose activated carbon (GK-HC), GK plus high-dose wheatgrass (GK-HW), GK plus high-dose dandelion (GK-HD) and GK plus high-dose corn stigma (GK-HS). Control Wistar (W0), Wistar plus medium-dose activated carbon (W-MC), Wistar plus medium-dose wheatgrass (W-MW), Wistar plus medium-dose dandelion (W-MD) and Wistar plus medium-dose corn stigma (W-MS) were also used to determine the treatment effects of drugs under healthy condition. Rats were housed in solid-bottom stainless steel cages with four per cage. During 30 days of the intervention, the rats were provided with free access to deionized water and received fresh drug administration via oral gavage once daily. Meanwhile, rats in the GK and W control group were administrated with deionized water. Blood glucose and weight of rats were measured on days 2, 9, 16, 23 and 30 before daily drug administration, and the fresh fecal samples were collected in sterile tubes for measurement of the gut microbiome and metabolites (Fig. S1). All feces were stored at −80 ^0^C until processing. At the end of the experiment, all rats were euthanized by cervical dislocation.

### Diet ingredients

The activated carbon and wheatgrass (*Triticum aestivum* L.) used in this study were obtained from Fujian Yuanli Activated Carbon Co., Ltd and Jiangsu Runhui Food Co., Ltd, respectively. Dandelion (*Taraxacum Officinale*) and corn stigma (*Stigma Maydis*) were provided by Hangzhou Lingyin Trading Co., Ltd and Kunming Xuanqing Biotechnology Co., Ltd, respectivelly. All of these drugs provided by producers were pulverized into 100-200 mesh particles. Corn starch was supplied by Nanjing Ganzhiyuan Sugar Co., Ltd. For further details on diet element composition, see Table S1.

The rats in the high dose group were administrated 1 g/kg of activated carbon, 0.15 g/kg of wheatgrass, 1 g/kg of dandelion and 0.15 g/kg of corn stigma, while the low dose group received 0.5, 0.07, 0.5, 0.07g/kg of the corresponding drugs. The healthy Wistar groups, on the other hand, were given 0.75, 0.1, 0.75, 0.1g/kg of the above drugs. Each drug was mixed with corn starch in a 1:1 ratio, and the drug-starch mixture were dissolved in 2.5 ml deionized water. The suspension was administered to each rat in the experiment groups by oral gavage for 30 consecutive days. The control groups were gavaged with an equal volume of deionized water, as previously mentioned. The suspension is freshly prepared every day, with the rest drug is stored in a dry environment until the second preparation.

### Measurements of fasting blood glucose (FBG)

After a 16-hour fasting, blood samples were taken from the tip of the tail vein on the 2nd, 9th, 16th, 23rd and 30th day for measurement of FBG using a glucometer and glucose strips (Sanocare Inc.).

### DNA extraction and 16S rRNA sequencing

After 0.1-mm glass bead beating for 5 min to ensure lysis of bacterial cells, total DNA was extracted from the fecal samples using QIAamp® Fast DNA Stool Mini Kit (Qiagen, Hilden, Germany) according to the manufacturer’s protocol. The DNA concentration was determined using Qubit, and the DNA samples were stored at −20 ^0^C until further use. The total DNA was used as a template for PCR amplification of the V3-V4 region of the 16S rRNA genes with universal primer sets 338F (5’-ACTCCTACGGGAGGCAGCAG-3’) and 806R (5’-GGACTACHVGGGTWTCTAAT-3’) (43).

PCR amplicon were purified and sequenced using an Illumina sequencing platform (MiSeq PE300, Meiji biological medicine Co., Ltd. China). Sequences were analyzed using the Quantitative Insights into Microbial Ecology (QIIME2, v2020.11). Amplicon sequence variants (ASVs) picking and taxonomy classification were performed based on sequence similarity by using DADA2 software with the SILVA v138 reference database. Alpha diversity was measured using Shannon and Chao1 diversity indices, beta diversity was indicated by calculating the weighted UniFrac dissimilarity matrix and visualized using principal coordinates analysis (PCoA). Linear discriminant analysis (LDA) effect size (LefSe) was applied to identify the most discriminant taxa among groups based on the feature table, with parameters set to default P value (α = 0.05) and LDA score of 3.0. PICRUSt2 was used to explore the potential functions in the gut microbiota, with predicted gene counts used to rarefy samples for further diversity analysis. The raw reads were deposited into the NCBI Sequence Read Archive (SRA) database under SRA accession number PRJNA936635.

### Untargeted metabolomic profiling

A total of 60 mg samples were extracted with 400 µL extraction solution (methanol:water, 4:1) for LC-MS/MS analysis. The extraction procedure included cryogenic grinding for 6min, ultrasonic extraction in a water bath for 30min, and incubation at −20 ^0^C for 30 min. After centrifugation at 13,000 g for 15min at 4 ^0^C, the supernatant was obtained for LC-MS/MS analysis of metabolites using a UHPLC-Q Exactive system (Thermo Fisher Scientific) equipped with the electrospray ionization (ESI) Turbo Ion-Spray interface and operated in both positive and negative ion mode (44,45). To assess the repeatability of the whole analysis process, all samples of equal volume were mixed to create a quality control sample (QC), which was inserted into a QC sample queue during instrument analysis. The Progenesis QI (WatersCorporation, Milford, USA) was used to process the raw data, resulting in a normalized data matrix comprising peak intensity, mass-to-charge ratio, and retention duration. Metabolite information was obtained from the metabolic public database (HMDB) (http://www.hmdb.ca/) and KEGG (https://www.genome.jp/kegg/).

The preprocessed data were analyzed using the Majorbio Cloud Platform (https://www.majorbio.com) (46). Partial Least Squares Discriminant Analysis (PLS-DA) was employed to evaluate the differences in metabolic profiles between groups. Metabolites with *P* < 0.05, VIP > 2 and a value of 1.5 FC (fold change) were selected as differential metabolites. Volcano plots were used to display filtered metabolites of interest. Metabolic pathway of these metabolites was selected based on the KEGG database. Procrustes analysis was conducted for both the metabolome and microbiome using the vegan R package. Correlations between gut microbial populations and metabolic parameters were summarized by Pearson correlation.

### Statistical analysis

Descriptive statistics in the study were calculated using Excel 2019. Unless stated otherwise, differences among groups were determined using one-way ANOVA with Tukey’s post hoc test and nonparametric Kruskal-Wallis tests in SPSS v26. *P* < 0.05 was considered statistically significant for all analyses (FDR adjusted). Alpha diversity was calculated using Past3. Pairwise PERMANOVA was used to test Weighted UniFrac distances and Bray-Curtis distances between microbiota communities and metabolites, using the vegan R package. Other graphics in this study were produced by Origin.

## Acknowledgements

This work was supported by the National Natural Science Foundation of China (42161134002) and the Alliance of International Science Organizations (Grant No. ANSO-CR-KP-2021-08).

Caixia Zhao: Project administration, Investigation, Resources, Data curation, Formal analysis, Software, Visualization, Writing – original draft. Yutong Wu: Project administration, Investigation, Resources, Methodology, Software. Jianqiang Su and Yin Wang: Project administration, Conceptualization, Supervision, Writing – review and editing, Funding acquisition.

## References

1. Cho N H, et al. 2018. Idf Diabetes Atlas: Global estimates of diabetes prevalence for 2017 and projections for 2045 [J]. Diabetes Research and Clinical Practice, 138: 271–281. 10.1016/j.diabres.2018.02.023.

2. Bellou V, et al. 2018. Risk factors for type 2 diabetes mellitus: An exposure-wide umbrella review of meta-analyses [J]. Plos One, 13(3): e0194127. 10.1371/journal.pone.0194127.

3. Walter J and Ley R. 2011. The human gut microbiome: ecology and recent evolutionary changes [J]. Annual Review of Microbiology, 65: 411–429. 10.1146/annurev-micro-090110-102830.

4. Deepa M, et al. 2017. Role of lifestyle factors in the epidemic of diabetes: lessons learnt from India [J]. European Journal of Clinical Nutrition, 71(7): 825–831. 10.1038/ejcn.2017.19.

5. Ross M C, et al. 2015. 16s gut community of the Cameron County Hispanic Cohort [J]. Microbiome, 3: 7. 10.1186/s40168-015-0072-y.

6. Proctor L M, et al. 2019. The Integrative Human Microbiome Project [J]. Nature, 569(7758): 641–648. 10.1038/s41586-019-1238-8.

7. Fan Yand Pedersen O. 2021. Gut microbiota in human metabolic health and disease [J]. Nature Reviews Microbiology, 19(1): 55–71. 10.1038/s41579-020-0433-9.

8. Vadder F D, et al. 2014. Microbiota-generated metabolites promote metabolic benefits via gut-brain neural circuits [J]. Cell, 156(1-2): 84–96. 10.1016/j.cell.2013.12.016.

9. Qin J, et al. 2012. A metagenome-wide association study of gut microbiota in type 2 diabetes [J]. Nature, 490(7418): 55–60. 10.1038/nature11450.

10. Zhao L, et al. 2018. Gut bacteria selectively promoted by dietary fibers alleviate type 2 diabetes [J]. Science, 359(6380): 1151–1156. 10.1126/science.aao577.

11. Hosomi K, et al. 2022. Oral administration of Blautia wexlerae ameliorates obesity and type 2 diabetes via metabolic remodeling of the gut microbiota [J]. Nature Communications, 13(1): 4477. 10.1038/s41467-022-32015-7.

12. Nguyen N K, et al. 2020. Gut microbiota modulation with long-chain corn bran arabinoxylan in adults with overweight and obesity is linked to an individualized temporal increase in fecal propionate [J]. Microbiome, 8(1): 118. 10.1186/s40168-020-00887-w.

13. Adhikary M, et al. 2021. Flavonoid-rich wheatgrass (Triticum aestivum L.) diet attenuates diabetes by modulating antioxidant genes in streptozotocin-induced diabetic rats [J]. Journal of Food Biochemistry, 45(4): e13643. 10.1111/jfbc.13643.

14. Gil B-S, et al. 2015. The Medical Use of Wheatgrass: Review of the Gap Between Basic and Clinical Applications [J]. Mini-Reviews in Medicinal Chemistry, 15(12): 10.2174/138955751512150731112836.

15. Kenny O, et al. 2015. Quantitative Uplc-Ms/Ms analysis of chlorogenic acid derivatives in antioxidant fractionates from dandelion (Taraxacum officinale) root [J]. 50(3): 766–773. 10.1111/ijfs.12668.

16. Cho S-Y, et al. 2002. Alternation of hepatic antioxidant enzyme activities and lipid profile in streptozotocin-induced diabetic rats by supplementation of dandelion water extract [J]. Clinica Chimica Acta: 10.1016/S0009-8981(01)00762-8.

17. Sabiu S, et al. 2016. Kinetics of alpha-amylase and alpha-glucosidase inhibitory potential of Zea mays Linnaeus (Poaceae), Stigma maydis aqueous extract: An in vitro assessment [J]. Journal of Ethnopharmacology, 183: 1–8. 10.1016/j.jep.2016.02.024.

18. Yadavalli T, et al. 2019. Drug-encapsulated carbon (Decon): A novel platform for enhanced drug delivery [J]. Science Advances, 5(8): 10.1126/sciadv.aax0780.

19. Larsen N, et al. 2010. Gut microbiota in human adults with type 2 diabetes differs from non-diabetic adults [J]. Plos One, 5(2): e9085. 10.1371/journal.pone.0009085.

20. Turnbaugh P J, et al. 2006. An obesity-associated gut microbiome with increased capacity for energy harvest [J]. Nature, 444(7122): 1027–1031. 10.1038/nature05414.

21. Hernández M A G, et al. 2019. The Short-Chain Fatty Acid Acetate in Body Weight Control and Insulin Sensitivity [J]. Nutrients, 11(8): 10.3390/nu11081943.

22. Zhao L. 2013. The gut microbiota and obesity: from correlation to causality [J]. Nature Reviews Microbiology, 11(9): 639–647. 10.1038/nrmicro3089.

23. Allcock G H, et al. 2001. Neutrophil accumulation induced by bacterial lipopolysaccharide: effects of dexamethasone and annexin 1 [J]. Clinical and Experimental Immunology, 123(1): 62–67. 10.1046/j.1365-2249.2001.01370.x.

24. Heaver S L, et al. 2018. Sphingolipids in host-microbial interactions [J]. Current Opinion in Microbiology, 43: 92–99. 10.1016/j.mib.2017.12.011.

25. Boini K M, et al. 2017. Sphingolipids in obesity and related complications [J]. Frontiers in Bioscience-Landmark: 10.2741/4474.

26. Johnson E L, et al. 2020. Sphingolipids produced by gut bacteria enter host metabolic pathways impacting ceramide levels [J]. Nature Communications, 11(1): 2471. 10.1038/s41467-020-16274-w.

27. Guo Q, et al. 2019. Hypoglycemic effects of polysaccharides from corn silk (Maydis stigma) and their beneficial roles via regulating the PI3k/Akt signaling pathway in L6 skeletal muscle myotubes [J]. International Journal of Biological Macromolecules, 121: 981–988. 10.1016/j.ijbiomac.2018.10.100.

28. Molinari R, et al. 2022. Polyphenols as modulators of pre-established gut microbiota dysbiosis: State-of-the-art [J]. Biofactors, 48(2): 255–273. 10.1002/biof.1772.

29. Tresserra-Rimbau A, et al. 2019. Associations between Dietary Polyphenols and Type 2 Diabetes in a Cross-Sectional Analysis of the Predimed-Plus Trial: Role of Body Mass Index and Sex [J]. Antioxidants (Basel), 8(11): 10.3390/antiox8110537.

30. Ahrens A P, et al. 2021. A Six-Day, Lifestyle-Based Immersion Program Mitigates Cardiovascular Risk Factors and Induces Shifts in Gut Microbiota, Specifically Lachnospiraceae, Ruminococcaceae, Faecalibacterium prausnitzii: A Pilot Study [J]. Nutrients, 13(10): 10.3390/nu13103459.

31. Meehan C J and Beiko R G. 2014. A phylogenomic view of ecological specialization in the Lachnospiraceae, a family of digestive tract-associated bacteria [J]. Genome Biology and Evolution, 6(3): 703–713. 10.1093/gbe/evu050.

32. Lin H, et al. 2016. Correlations of Fecal Metabonomic and Microbiomic Changes Induced by High-fat Diet in the Pre-Obesity State [J]. Scientific Reports, 6: 21618. 10.1038/srep21618.

33. Zierer J, et al. 2018. The fecal metabolome as a functional readout of the gut microbiome [J]. Nature Genetics, 50(6): 790–795. 10.1038/s41588-018-0135-7.

34. Yang S J, et al. 2018. Serum metabolite profile associated with incident type 2 diabetes in Koreans: findings from the Korean Genome and Epidemiology Study [J]. Scientific Reports, 8(1): 8207. 10.1038/s41598-018-26320-9.

35. Yan L, et al. 2021. Discovery of lipid profiles of type 2 diabetes associated with hyperlipidemia using untargeted Uplc Q-Tof/Ms-based lipidomics approach [J]. Clinica Chimica Acta, 520: 53–62. 10.1016/j.cca.2021.05.031.

36. Cruickshank-Quinn C, et al. 2017. Metabolomic similarities between bronchoalveolar lavage fluid and plasma in humans and mice [J]. Scientific Reports, 7(1): 5108. 10.1038/s41598-017-05374-1.

37. Kuehbacher T, et al. 2008. Intestinal TM7 bacterial phylogenies in active inflammatory bowel disease [J]. Journal of Medical Microbiology, 57(Pt 12): 1569–1576. 10.1099/jmm.0.47719-0.

38. Zhu F-C, et al. 2020. Genomic Characterization of a Novel Tenericutes Bacterium from Deep-Sea Holothurian Intestine [J]. Microorganisms, 8(12): 10.3390/microorganisms8121874.

39. Zeuthen L H, et al. 2006. Lactic acid bacteria inducing a weak interleukin-12 and tumor necrosis factor alpha response in human dendritic cells inhibit strongly stimulating lactic acid bacteria but act synergistically with gram-negative bacteria [J]. Clinical And Vaccine Immunology, 13: 10.1128/Cvi.13.3.2006.365-375.

40. Wang Y, et al. 2021. The protective mechanism of a debranched corn starch/konjac glucomannan composite against dyslipidemia and gut microbiota in high-fat-diet induced type 2 diabetes [J]. Food Function, 12(19): 9273–9285. 10.1039/d1fo01233a.

41. Duncan S H, et al. 2009. The role of ph in determining the species composition of the human colonic microbiota [J]. Environmental Microbiology, 11(8): 2112–2122. 10.1111/j.1462-2920.2009.01931.x.

42. Walter J and Ley R. 2011. The Human Gut Microbiome: Ecology and Recent Evolutionary Changes [J]. Annual Review of Microbiology, 65(1): 411–429. 10.1146/annurev-micro-090110-102830.

43. Huang H, et al. 2022. Metabonomics combined with 16s rrna sequencing to elucidate the hypoglycemic effect of dietary fiber from tea residues [J]. Food Res Int, 155: 111122. 10.1016/j.foodres.2022.111122.

44. Zhao H, et al. 2022. A pilot exploration of multi-omics research of gut microbiome in major depressive disorders [J]. Transl Psychiatry, 12(1): 8. 10.1038/s41398-021-01769-x.

45. Want E J, et al. 2010. Global metabolic profiling procedures for urine using Uplc–Ms [J]. Nature Protocols, 5: 1005–1018. 10.1038/nprot.2010.50.

46. Ren Y, et al. 2022. Majorbio Cloud: A one-stop, comprehensive bioinformatic platform for multiomics analyses [J]. 1(2): e12. 10.1002/imt2.12.

